# Deep Neural Networks Point to Mid-level Complexity of Rodent Object Vision

**DOI:** 10.1101/2020.02.08.940189

**Authors:** Kasper Vinken, Hans Op de Beeck

## Abstract

In the last two decades rodents have been on the rise as a dominant model for visual neuroscience. This is particularly true for earlier levels of information processing, but high-profile papers have suggested that also higher levels of processing such as invariant object recognition occur in rodents. Here we provide a quantitative and comprehensive assessment of this claim by comparing a wide range of rodent behavioral and neural data with convolutional deep neural networks. These networks have been shown to capture the richness of information processing in primates through a succession of convolutional and fully connected layers. We find that rodent object vision can be captured using low to mid-level convolutional layers only, without any convincing evidence for the need of higher layers known to simulate complex object recognition in primates. Our approach also reveals surprising insights on assumptions made before, for example, that the best performing animals would be the ones using the most complex representations – which we show to likely be incorrect. Our findings suggest a road ahead for further studies aiming at quantifying and establishing the richness of representations underlying information processing in animal models at large.

## 1 Introduction

Up to one decade ago, macaque monkeys were uncontested as the primary animal model for vision research targeting information processing at the cortical level. Since then, studies on rodents have become increasingly popular, in particular because developments in neurotechnology such as cell-level imaging and optogenetics capitalized upon the existence of genetic models. A major question emerging from this evolution is how far we can take rodents as a model for the more complex aspects of visual information processing.

To answer this question, a series of high-profile studies have documented the ability of rats to perform seemingly complex object recognition tasks. In these tasks, rats show the ability to recognize objects despite various transformations in viewing conditions such as position, size, and viewpoint. In the first landmark study, Zoccolan and colleagues (2009) trained rats to discriminate two computer-graphics rendered objects under various changes in size and rotation, and showed that the animals could successfully generalize to novel combinations of these transformations. Several studies have subsequently expanded on this work using similar stimuli (Tafazoli et al., 2012; Alemi-Neissi et al., 2013; Rosselli et al., 2015), and recently it was found that the success of rats in these tasks depends on the complexity of their strategy (Djurdjevic et al., 2018). In a study using natural stimuli, rats could generalize learned category rules to new, unseen category exemplars that differ from trained exemplars in complex ways (Vinken et al., 2014). Neurophysiological recordings have revealed neural responses in lateral visual areas that might underlie these behavioral abilities (Vermaercke et al., 2014; Tafazoli et al., 2017). To cite Zoccolan (2015), “The picture emerging from this survey is very encouraging with regard to the possibility of using rats as complementary models to monkeys in the study of higher-level vision.”

However, this hypothesis is not yet proven scientifically. Up to now, the predictions were not specific enough. Already from the first landmark study by Zoccolan et al. (2009), it has been argued that the behavior is unlikely to be based upon representations at the level of primary visual cortex (V1). However, in primates there is a multi-step progression of complexity in representations beyond V1. The first processing stages are gradually building on top of the V1 representations, still containing properties such as local receptive fields and a retinotopic organization. In the latest stages, other, more complex representational properties start to dominate, from which object and category information can be easily read out (Dicarlo et al., 2012). If not based on a V1-like representation, then would rodent behavior in object vision tasks be supported by a V2-type representation or, at the other end of the range of possibilities, an object- and category-selective representation?

Here we demonstrate the added value of a computational framework to address this issue. Recently, a family of computational models has emerged in the form of convolutional deep neural networks (DNNs) that allow to simulate this hierarchical information processing. When trained on object recognition, DNNs show interesting commonalities with the primate ventral stream, with a progression of representations that is surprisingly similar to what is seen in monkeys and humans for brief stimulus presentations (Cadieu et al., 2014; Yamins et al., 2014; Güçlü and van Gerven, 2015; Kalfas et al., 2017, 2018; Pospisil et al., 2018; Bashivan et al., 2019), as such capturing important aspects of object recognition and perceived shape similarity (Yamins et al., 2014; Kubilius et al., 2016; Kalfas et al., 2018). The architecture of these computational models is composed of series of convolutional layers that perform local filtering operations, followed by fully connected layers, which gradually transforms pixel level inputs into a high-level representational space where object categories are linearly separable. Importantly, these models are image computable and can be readily used to extract an internal representation from each of these layers for any visual stimulus set.

Here we use these DNNs to formalize the level of complexity that underlies the data obtained from rats in object recognition tasks, focusing upon the most high profile studies suggesting a high level of invariance and complexity. On top of the outputs of each layer of the DNNs, we trained a linear classifier on the tasks and stimuli that were used in these studies, and we check which layers are needed to capture the rat data. The findings are very consistent across datasets: the data from earlier studies meant to probe high-level object vision in rodents can be explained by low to mid-level convolutional representations that fall short of the complexity of representations that underlie object recognition in primates. Surprisingly, crucial aspects of some of the most high profile studies are in fact captured by the first convolutional layer operating on pixel values. This was not the case for an object recognition task with natural videos, which required multiple convolutional and pooling operations. These findings shall be an important benchmark for future studies aiming to develop animal models, rodents or other species, for human vision.

## 2 Results

We focus on three landmark studies: the first showed that rats can learn to discriminate two objects invariant to changes in size and azimuth-rotation (Zoccolan et al., 2009), the second found that rats employ different object recognition strategies that seem to vary in complexity (Djurdjevic et al., 2018), the third showed that rats are capable of ordinate-level categorization of natural videos (Vinken et al., 2014).

### 2.1 Any DNN layer can account for size and rotation tolerance in a task with two rendered objects

We started by assessing the complexity of processing required in the first landmark paper that reported evidence for invariant object recognition in rats (Zoccolan et al., 2009). In this study, rats were first trained to discriminate two different objects and to tolerate variations in size and azimuth-rotation. At each trial, one object was presented and the rat had to indicate the object identity by licking either a left or right feeding tube. After training, the rats were tested on the full stimulus set, which included novel combinations of size and azimuth-rotation (Fig. 1**a,b**).

**Fig. 1.**
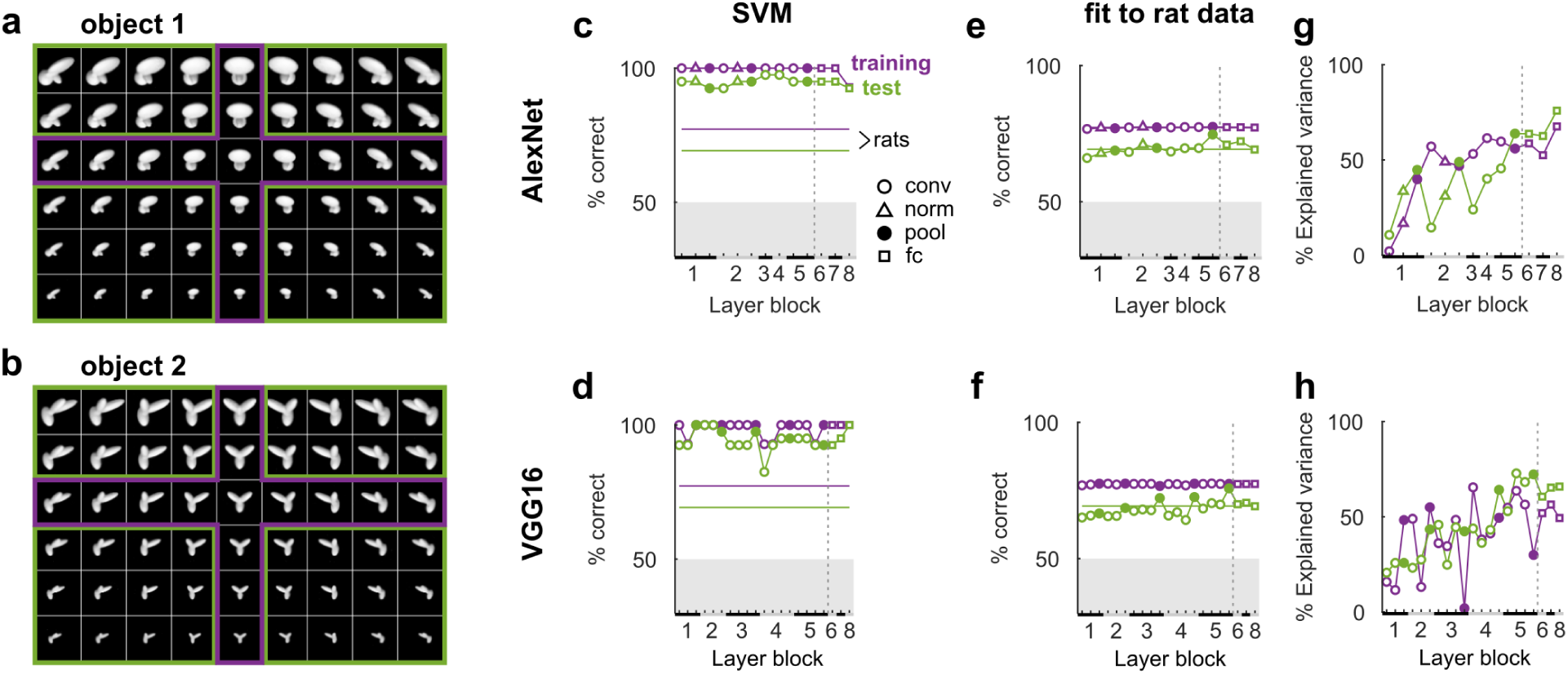
The first DNN layers account for generalization performance across a range of transformations, but higher layers explain more variance in transformation-level behavioral patterns in Zoccolan et al. (2009). **a,b**, the full sets of size and azimuth-rotation combinations of the two objects used in the behavioral task. Rats were first trained on a subset of these transformations (purple) and subsequently asked to generalize to novel combinations (green). **c,d**, average percentage correct discrimination of object 1 and object 2 by a linear classifier (SVM) operating on the outputs of each layer in AlexNet (**c**) and VGG16 (**d**). The classifier was first trained on the same object transformations as the rats, after which performance was evaluated on these trained transformations (purple) as well as the untrained transformations (green). Horizontal lines indicate the average behavioral performance across rats reported by Zoccolan et al. (2009). Black and grey bars on the X-axis indicate layer blocks and markers indicate layer types (see legend insert); the division between convolutional and fully connected layer blocks is indicated by a dashed line. **e,f**, average percentage correct discrimination estimated by a logistic regression mapping (Methods, Computational modeling, Behavioral tasks) for AlexNet (**e**) and VGG16 (**f**). **g,h**, transformation-level explained variance in percentages correct based on the mapping in **e,f**, respectively. Conventions for **e-h** are the same as in **c,d**.

In the original experiment, the rats generalized remarkably well to the object transformations they had never seen before. We trained DNN-based models using layers from AlexNet and VGG16 on the same task and found that this level of generalization turned out to be surprisingly easy for even the earliest layers: the models based on the first convolutional layer of both AlexNet and VGG16 already achieved near perfect generalization performance on the test set (Fig. 1**c,d**). Thus, very little processing is required to explain a high level of generalization performance from the trained to untrained object transformations.

In addition, we tested whether these models also explain transformation-level differences in behavioral performance between exemplar images. Because generalization performance of the DNNs was at ceiling for most object transformations, we used a logistic regression approach to map distances to the SVM decision boundary onto the average rat performances for each object transformation, based on the rat training data only (Methods, Computational modeling, Behavioral tasks). Next, we used this behavioral mapping to predict generalization performances for each of the novel (green) object transformations. On average, for each layer the behavioral mapping generalized well to the test set and was able to capture the difference in average training-test performance of the rats (Fig. 1**e,f**). We then calculated the transformation-level variance in behavioral performance that was explained by the DNN models. These results show that the first convolutional layer only explains a limited amount of transformation-level variance, which then sharply increases and slowly reaches a maximum in the highest convolutional layers (Fig. 1**g,h**). Thus, whereas representations in the first layer of AlexNet and VGG16 can already explain successful generalization to untrained object transformations, higher convolutional layers get increasingly better at explaining transformation-level differences in behavioral performance of rats. None of the networks shows a clear further improvement from incorporating the fully connected layers.

### 2.2 Earliest DNN layers can account for stimulus-level performance patterns across rendered objects and sizes, also for the best performing rats

To further investigate how well the DNN models can explain object and transformation-level differences in the behavioral performance of rats, we turned to a recent study by Djurdjevic et al. (2018). In this study, rats were trained to discriminate a reference object from 11 distractor objects at different sizes (Fig. 2**a**), with a similar experimental paradigm as in Zoccolan et al. (2009). The goal of the study was to use the variable discrimination performance observed across object conditions to infer the complexity of the rats’ perceptual strategy. The authors found that there were “good performers”, which performed above chance for the most challenging distractors, and “poorer performers”, which performed below chance for the most challenging distractors.

**Fig. 2.**
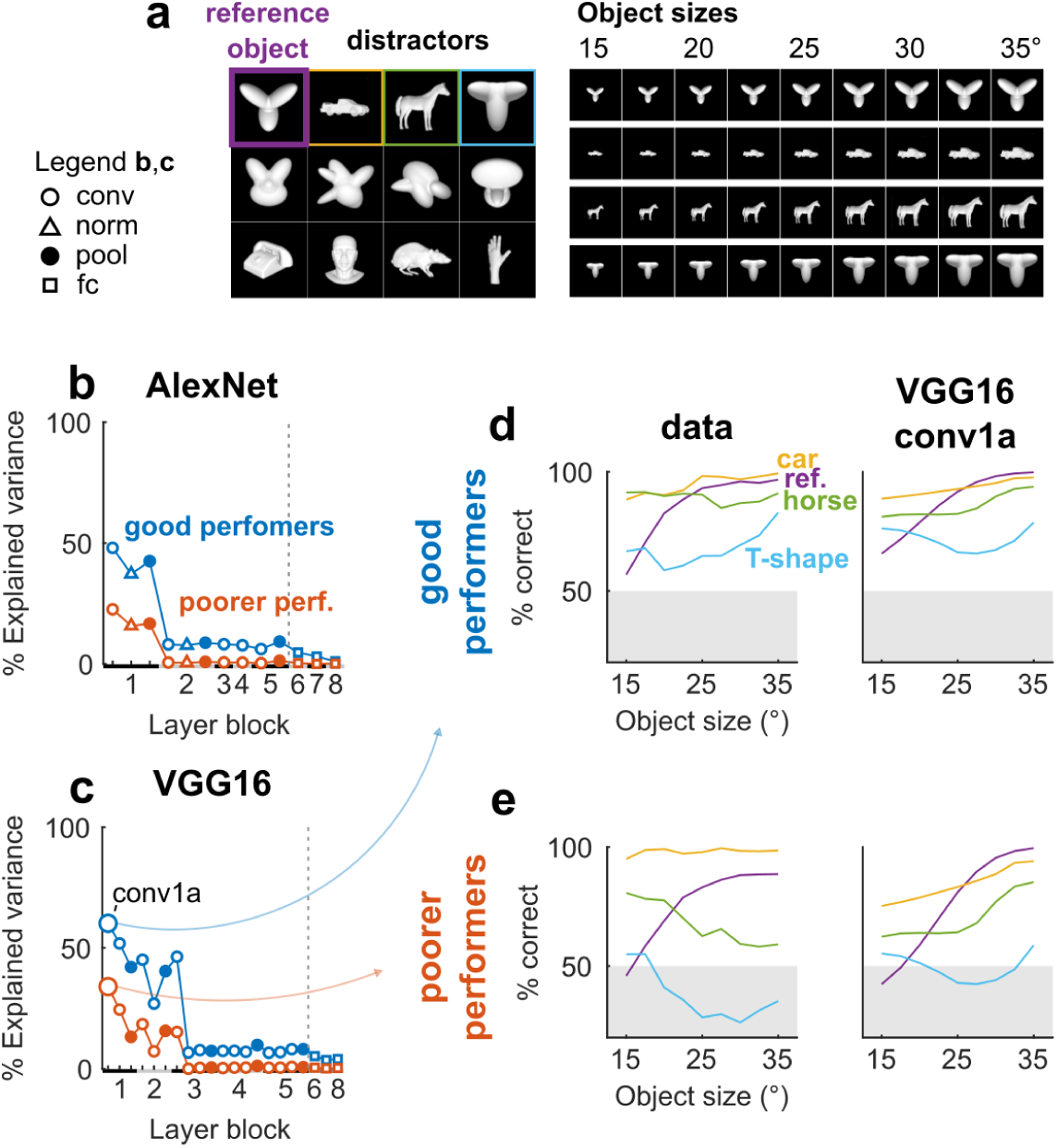
Discrimination performance patterns for objects from Djurdjevic et al. (2018) map onto earliest DNN layers. **a**, left: the reference object (purple) and 11 distractor objects (rest) that rats were trained to discriminate in the behavioral task. The 3 example distractors of Figure 1 in Djurdjevic et al. (2018) are highlighted in yellow, green, and blue. Right: in a later phase of the experiment, the rats were trained to tolerate size changes in all objects from 15° to 35° of visual angle (here only shown for the subset of 4 objects indicated in color on the left). **b**, percentage of variance in average discrimination performance of good performers (blue) and poorer performers (red), explained by the distance to the SVM boundary for each layer in AlexNet (Methods, Computational modeling, Behavioral tasks). This time a bias term (intercept) was included to capture the difference between good and poorer performers. The behavioral mapping was estimated using the data reported for the 4 example stimuli in Figure 1 of Djurdjevic et al. (2018), and the explained variance was calculated for the same data. Black and grey bars on the X-axis indicate layer blocks and markers indicate layer types (see legend insert); the division between convolutional and fully connected layer blocks is indicated by a dashed line. **c**, same as **b**, but for VGG16. Conv1a (indicated by the larger markers) explains the most variance. **d**, left: average discrimination performances of good performers, as a function of object size, for the reference object and the 3 example distractors (these are the data used for the behavioral mapping). Right: average discrimination performances of good performers, predicted from the behavioral mapping using the model that explained the most variance in **b** and **c** (i.e. conv1a in VGG16). **e**, same as **d**, but for poorer performers.

We calculated how much of the object and size-level variance in behavioral performance was explained by the DNN models, using the data that was displayed in Figure 1 of Djurdjevic et al. (2018). For both good and poorer performers, the percentage explained variance was highest for the first convolutional layer, and higher for VGG16 than AlexNet (Fig. 2**b-e**). Surprisingly, in all cases the percentage explained variance was higher for good performers than for poorer performers and this difference was largest for earliest layers, in contrast with the idea that good performers relied on more advanced processing of shape information (Djurdjevic et al., 2018). In sum, object and size-level differences in behavioral performance of rats were best explained by the layer with the least amount of processing, and better for the good performers, suggesting that the rats’ perceptual strategies could have relied on low-level visual representations.

### 2.3 Mid-level DNN layers can account for natural video categorization behavior

Up to this point, we have only discussed studies that used a small number of computer-graphics renderings of abstract and more naturalistic objects. However, rats have also been shown to be able to learn category rules from more complex natural videos that generalize to novel category exemplars (Vinken et al., 2014). In this study, rats were trained in a visual water maze (Prusky et al., 2000) to classify videos featuring a rat (target category), from phase scrambled versions of the target videos and target-matched natural distractor videos featuring various moving objects. In each trial the target and distractor were presented simultaneously, with one video on the left and the other on the right. The rats were first trained on a training set of 15 videos (Fig. 3**a**), initially with a fixed target-distractor pairing, followed by a phase where all possible target-distractor combinations were presented. In the subsequent test phase the rats were probed with 40 novel videos with a fixed target-distractor pairing (Fig. 3**b**), without negative feedback for incorrect trials.

**Fig. 3.**
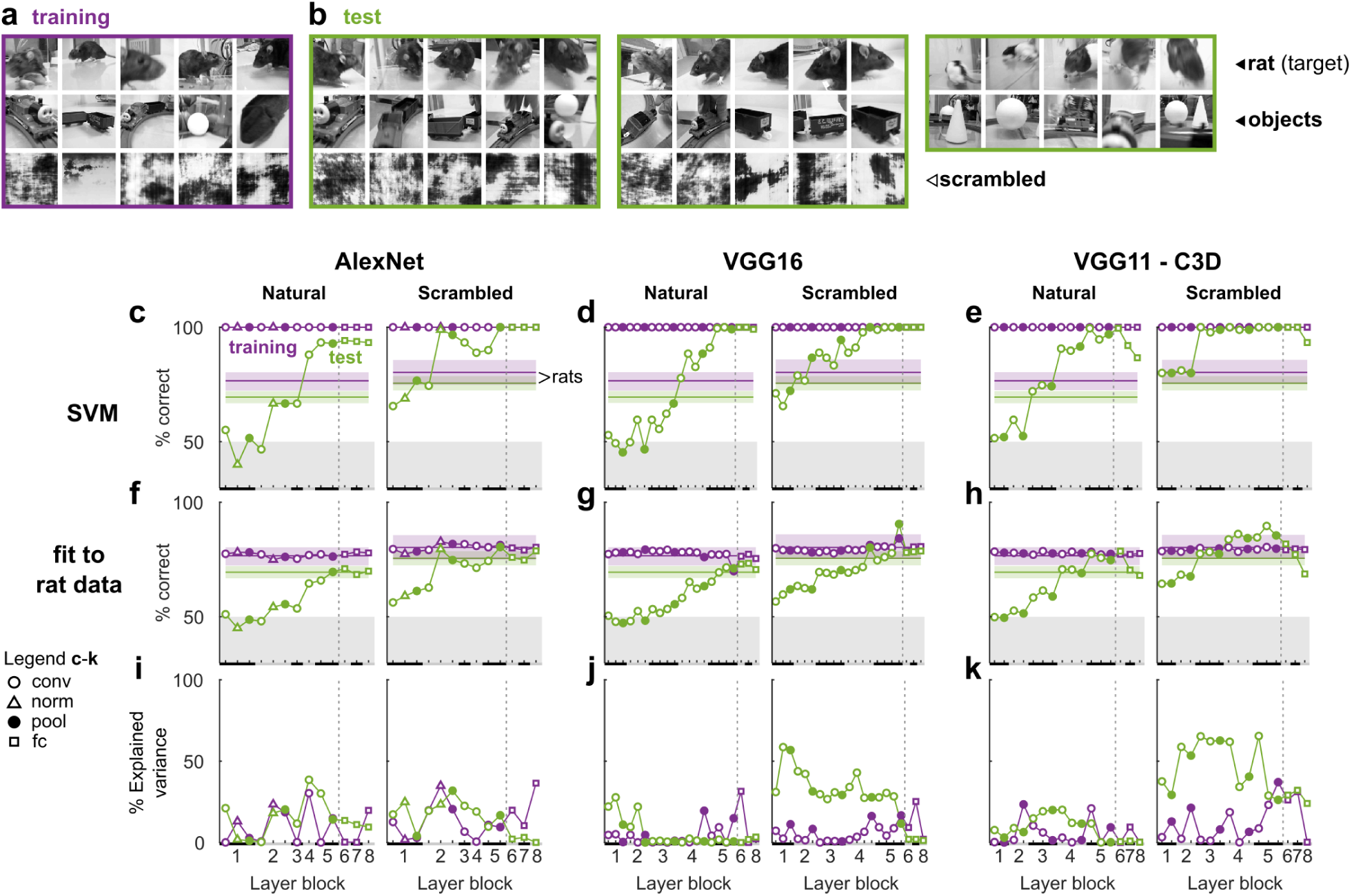
Mid-level convolutional layers are required for generalization in a rat versus non-rat categorization task of natural videos in Vinken et al. (2014). **a,b**, single frames of all the training (**a**) and test (**b**) videos that rats were asked to classify in the behavioral task. Each rat was trained a subset of 15 videos (purple), and tested for generalization with 40 novel videos (green). Ten test videos (the natural videos in the first green rectangle) were modified to further probe the rats, for example by reducing playback speed to 25% or equalizing average pixel values in the lower-half of the videos (see Vinken et al., 2014). **c-e**, average percentage correct classification of rat versus non-rat frame bin pairs by the classifier operating on the outputs of each layer in AlexNet, VGG16, and VGG11-C3D and for natural and scrambled distractors separately. Performance is evaluated on the training set (purple; all 50 target-distractor combinations) as well as those of the test set (green; 25 tested target-distractor pairs, with 25% playback and pixel value modifications for 5 pairs). Black and grey bars on the X-axis indicate layer blocks and markers indicate layer types (see legend insert); the division between convolutional and fully connected layer blocks is indicated by a dashed line. Horizontal lines indicate the average behavioral performance across rats and error bounds are 95% bias corrected accelerated bootstrap confidence intervals. **f-h**, average percentage correct classification of rat versus non-rat stimulus pairs estimated by a logistic regression mapping (Methods, Computational modeling, Behavioral tasks) onto to the actual rat performances for each stimulus pair. The parameters for the behavioral mapping were fit on the video pairs and data of the training set only. Conventions are the same as in **c-e. i-j**, target-distractor-level explained variance in performance based on the behavioral mapping. Conventions are the same as in **c-e**.

The rats were able to generalize well to the novel videos, independently of temporal information or local luminance cues (although local luminance did explain some of the stimulus-level response variance; Vinken et al., 2014). To test whether this generalization could be explained by other low-level visual information, we trained DNN-based models using layers from AlexNet, VGG16, and VGG11-C3D on the same two alternative forced choice task (Methods, Computational modeling, Behavioral tasks). VGG11-C3D is a convolutional neural network with 3D spatio-temporal filters (16-frame temporal bins) and thus able to encode temporal features (Tran et al., 2014). For AlexNet and VGG16 we averaged outputs across frames for each 16-frame bin. All three networks performed very similarly: for scrambled distractors the models based on the first layers already performed almost at the behavioral level of rats, but for natural distractors they were at chance and did not exceed rat-level performance until conv4 of AlexNet, conv4a for VGG16, and conv3a for VGG11-C3D (Fig. 3**c-e**).

We used logistic regression again to map differences in distances to the SVM decision boundary onto the average rat performances for each target-distractor combination of the training set and then used this behavioral mapping to predict test set generalization performances. On average, this behavioral mapping matched the observed gap between rat training and test set performance for even higher layers (Fig. 3**f-h**). For scrambled distractors, a relatively high target-distractor-level variance in behavioral performance was explained by the DNN models based on earlier layers to mid-level layers in the VGG architecture networks. However, for natural distractors the explained variance was generally low and highly variable across successive layers, making it harder to interpret (Fig. 3**i-k**).

Together, these results suggest that, in terms of DNNs trained on object recognition, a mid-level representation based on several convolutional and pooling operations is required to explain the observed generalization to novel category exemplars in a classification task of natural videos, while a modest amount of target-distractor-level variance is explained by early to mid-level layers. Again there is no evidence of an added benefit of fully connected layers.

### 2.4 Representations of natural and scrambled videos in the rat lateral stream change in parallel with representations in DNNs

DNN based models suggest that several early and intermediate layers of hierarchical processing in an object recognition model are required for a representation that can support natural video categorization. The rodent cortex houses a complex network of specialized higher-order visual areas (Glickfeld and Olsen, 2017), but is there any evidence for such a representation in the rat brain? A likely candidate pathway is the “lateral visual stream”, which anatomically resembles the primate ventral visual stream (Wang et al., 2012) and shows several functional properties thought to be typical of an object recognition pathway, such as increased tolerance for changes in position (Vermaercke et al., 2014), size, rotation, and illumination (Tafazoli et al., 2017).

Previously, we investigated neural representations of natural and scrambled videos along the rat lateral stream (Vinken et al., 2016). In this experiment, we presented the natural videos of the training set in Fig. 3**a** and their scrambled counterparts to awake, passively watching rats which were never trained with these videos. We recorded single and multi unit spiking activity in primary visual cortex (V1), a middle laterointermediate area (LI), and the most lateral temporal occipital cortex (TO). We found evidence for an increased dissociation of natural and scrambled videos from V1 to TO, but not for a categorical representation (Vinken et al., 2016).

Here, we investigated the similarity between neural representations of these videos in the rat lateral stream and the DNN representations in each layer of each model. We calculated representational dissimilarity matrices (RDMs) based on the neural responses per 16-frame time bin (Fig. 4**a-c**), and compared these with RDMs based on DNN layer activity (Fig. 4**d-f**). Visual inspection reveals that, similar to the neural RDMs, the DNN layer RDMs show a progression towards an increased dissociation between natural and scrambled videos. However, unlike the neural data, the RDMs of the last fc8 layers suggest a category-like representation differentiating the rat from the non-rat videos (see checkered pattern in the left-upper quadrant).

**Fig. 4.**
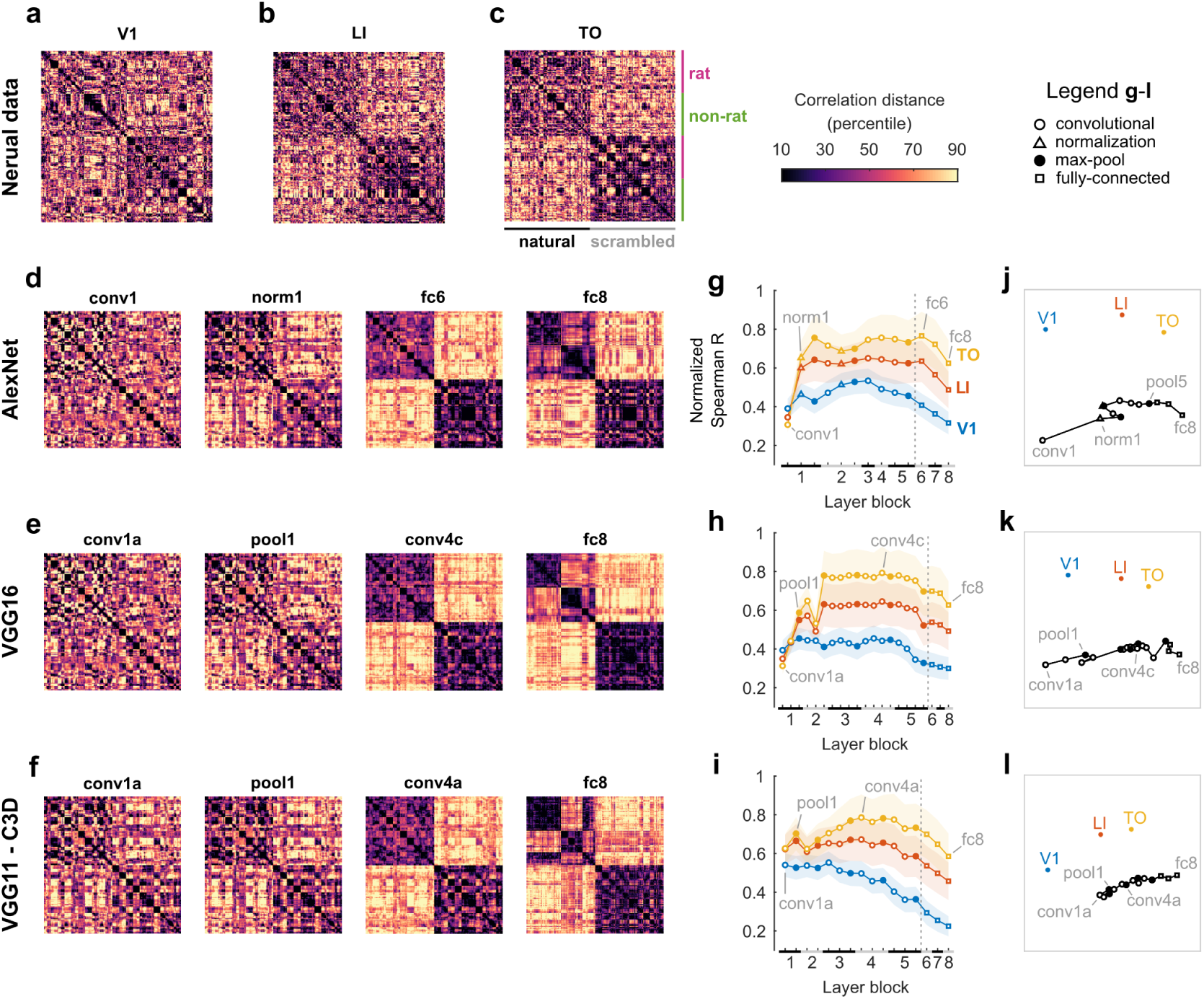
Neural representational dissimilarity matrices for natural and scrambled videos per lateral stream area change in parallel with DNN layer representations. **a-c**, neural RDMs for V1, LI, and TO data. Rows and columns correspond to 16-frame bins: nine per video, with first the five rat videos, then the five non-rat videos, and finally the scrambled versions of the natural videos in the same order. The color scale corresponds to percentiles of each RDM’s Pearson correlation distances (excluding the diagonal values). **d-f**, artificial neural network RDMs (Pearson correlation distance) for 4 layers of AlexNet, VGG16, and VGG11-C3D: the first layer, the earliest layer for which the RDM corresponds better to extra-striate (LI/TO) data, the layer with the best normalized correlation with neural data (TO in all three cases), and the last layer (fc8). **g-i**, Spearman correlations between each artificial neural network layer RDM and each neural RDM (calculated using above diagonal elements only), normalized by each area’s noise ceiling (V1 in blue, LI in red, TO in yellow). Black and grey bars on the X-axis indicate layer blocks and markers indicate layer types (see legend insert); the division between convolutional and fully connected layer blocks is indicated by a dashed line. Grey text labels indicate the DNN RDMs shown in **d-f**. Error bounds are 95% confidence intervals calculated using Jackknife standard error estimates. **j-l**, two-dimensional representation of similarities between neural and artificial neural network RDMs, derived from applying non-metric multidimensional scaling on Spearman correlation distances between RDMs. Each marker corresponds to an RDM and similar RDMs are plotted closer together. Text labels indicate the neural RDMs and the DNN RDMs shown in **d-f**.

A further quantitative comparison between each neural and DNN layer RDM showed that the similarity between neural and DNN layer representations increases for LI and TO in later layers, well beyond the similarity for V1 data (non overlapping 95% confidence intervals, Fig. 4**g-i**). When we visualized all between-RDM similarities using multidimensional scaling, the plots suggested a progression from V1 to TO parallel to the progression across successive DNN layers (Fig. 4**j-l**). The representational similarity was consistently higher for TO, peaking between pool1 and fc6 for AlexNet (Fig. 4**g**), pool2 and conv5a for VGG16 (Fig. 4**h**), and pool3 and conv5a for AlexNet (Fig. 4**i**).

Overall, these results suggest that the neural representations of the videos in the most lateral visual area TO correspond best to the mid-level representations in DNNs trained on object recognition.

## 3 Discussion

We formally investigated the level of processing required to explain behavioral performance in several landmark studies on rodent visual object recognition. Using convolutional neural network architectures trained on object recognition, we assessed at which stage of processing each neural network can solve the task. This computational approach provides a generally consistent picture of the representations that underlie the behavior of rats. All tasks could be captured by convolutional layers only, even for the most challenging task which required later convolutional layers (Fig. 3). This picture is much more precise than in earlier papers, which generally relied on the assumption that rats could not have generalized across variations in appearance and identity preserving transformations using trivial strategies; an assumption that up until now had not been tested scientifically.

Besides the general performance, there were interesting commonalities between behavior and neural network representations at a more fine-grained level. Despite the ability of the first convolutional layer to generalize across different sizes and rotations in Zoccolan et al. (2009), later convolutional layers could explain more transformation-level variance in performance (Fig. 1). On the other hand, performance differences between objects and sizes in Djurdjevic et al. (2018) were best explained by the earliest convolutional layers, also for the best performing rats (Fig. 2). Finally, consistent with our finding in Fig. 3 that later convolutional layers capture the behavioral capacities shown by Vinken et al. (2014), later convolutional layers matched the representational geometry increasingly better for extrastriate areas in the rat visual cortex (Fig. 4).

While our comparisons with DNNs show that rodent object vision can be explained by convolutional layers, in primates fully connected layers of the same or similar network architectures capture perceived shape similarity better than convolutional layers (Kubilius et al., 2016; Kalfas et al., 2018; Bracci et al., 2019). This suggests that the rodent visual system is capable of a mid-level complexity of object processing which is markedly less complex than primates. For the similarity between neural representations in primate inferotemporal cortex and DNN layers the story is less consistent: in some cases it does peak at the fully connected layers (Khaligh-Razavi and Kriegeskorte, 2014), whereas in other studies it peaks at the last convolutional layer (Kalfas et al., 2018; O’Connell et al., 2019; Güçlü and van Gerven, 2015; Grossman et al., 2019), suggesting that there might be a dependence on the particular stimulus set. This shows the road ahead for future studies on comparing behavior and neurophysiology between animal species: preferably the exact same stimuli and tasks would be used, designed to be able to differentiate among representations and strategies of varying complexity. Currently no such data are available.

Our results provide important qualifications for assumptions about the stimulus set or task complexity that are made by researchers. First, in Zoccolan et al. (2009), the overall performance level, is surprisingly easy to explain based on activations in the first convolutional layer. Instead, a more in-depth look at the behavioral performance patterns across object transformations turned out to be more relevant for showing at least mid-level processing. Second, in Djurdjevic et al. (2018), behavioral patterns for good performers are actually more consistent with early convolutional representations, contrary to intuitive assumption of the authors that these animals must rely on a complex strategy. Third, a paradigm where a only single stimulus is presented on every trial (as in Zoccolan et al., 2009; Djurdjevic et al., 2018) has been suggested (Zoccolan, 2015) to be better to probe more complex strategies compared to a two-alternative forced choice task where the target and distractor can be directly compared Vinken et al. (2014). However, here we show that, if anything, the latter paradigm can provide evidence for representations that are at least as complex as the paradigm used by Zoccolan et al. (2009).

These considerations are of critical importance for future research on rodent vision. For example, one of the most large-scale initiatives to investigate the visual system in rodents is headed by the Allen Institute, where the most recent efforts yielded nearly 100,000 recorded neurons (Siegle et al., 2019; De Vries et al., 2018). The interpretation of the neural data is greatly facilitated when behavioral read-outs are available. The current study points to two challenges in this respect. First, we need a computational approach to validate assumptions about the complexity of the visual tasks and the strategies used for task performance. Second, prior to starting data collection, a computational approach would be highly valuable at the stage of deciding which stimuli to use. Even for the large scale neural recordings (De Vries et al., 2018), the bottleneck for comprehensively characterizing higher-level visual processing could be in the design of the stimulus set rather than the number of recorded neurons.

More generally, the consideration about researcher assumptions is not limited to rodent studies. The vast majority of monkey and human work lacks a computational framework to evaluate assumptions made by the researchers about their paradigm, stimulus set, or resulting data. Yet, as we demonstrate here, perfectly reasonable assumptions about task difficulty or interpretations of patterns in the data can turn out to be incorrect when comprehensively tested using computational models. Therefore, whenever applicable, the use of tried-and-tested computational models such as DNNs should prove invaluable as a standard methodological practice for the design of experiments on high-level (visual) cognition as well as for a precise and correct interpretation of the results.

Finally, it is important to emphasize that the fact that we have not found any evidence for truly high-level visual object recognition behavior does not imply that it is not there or cannot be there. While our results are inconsistent with the idea that rodents are nocturnal animals that rely more extensively on whisker touch and smell when exploring their environment, it is also possible that none of the studies discussed here really pushed the limits of rodent object vision. A road ahead for addressing this question is to use DNNs to construct stimulus sets and design paradigms that explore the boundaries of rodent vision by getting the best out of them. This will likely also include considerations about the ecological validity of the tasks from the perspective of rodents (Cox, 2014; Vinken et al., 2017; Cadena et al., 2019). Another interesting avenue is the use of paradigms that elucidate the strategies used by rats by means of experimental manipulations that also made the task more challenging and exclude some of the simplest pixel-based strategies. For example, Alemi-Neissi et al. (2013) presented stimuli covered by masks that randomly occluded parts of the objects, which effectively excludes the possibility of the animals using one highly specific local-luminance based strategy. In addition, in a second part of their study, Djurdjevic et al. (2018) presented random variations of the reference image to infer perceptual templates, showing that a template-matching model using only one fixed perceptual template could not account for the animal’s performance across image manipulations such as changes in object size. Given the difficulty to relate these strategies to the complexity of the underlying representations, it will be interesting to combine a more computational DNN approach with these template paradigms.

In summary, we used convolutional deep neural networks for a comprehensive and quantitative assessment of the level of processing required to explain rodent visual object recognition. A combination of behavioral and neural results reveals a mid-level complexity of visual processing, consistent with the idea of a visual system that is reasonably advanced but not the primary modality. The main conclusion can be phrased in different ways, depending on whether one takes a glass half-full or half-empty perspective. At one hand, our findings confirm that rodent visual task performance is non-trivial, displays a certain degree of invariance, and requires multi-layer networks to be simulated. At the other hand, the behavioral performance as well as underlying neural representations point to underlying representations of a limited complexity relative to primate vision.

## 4 Methods

### 4.1 Behavior

#### 4.1.1 Data and stimulus extraction

We extracted the data and stimuli from the manuscripts of Zoccolan et al. (2009) and Djurdjevic et al. (2018). The object images were directly copied from Fig. 2A and Figure 1A, respectively. The average rat performances were copied from the numbers in Fig. 2B of (Zoccolan et al., 2009, *N* = 6 rats), and extracted from Figure 1C of Djurdjevic et al. (2018) using WebPlotDigitizer 4.2 (https://apps.automeris.io/wpd/; *N* = 6 rats). For methodological details about the experimental setup and tasks, we refer to the original papers.

#### 4.1.2 Natural video categorization

We used the original movies from Vinken et al. (2014), and extracted the behavioral data (*N* = 5 rats) from the data files of that study. For methodological details about the experimental setup and the task, we refer to the original paper.

### 4.2 Neurophysiology

#### 4.2.1 Data

The neurophysiological data are from a study published previously in (Vinken et al., 2016, *N* = 7 rats) and consist of single cell and multi unit responses to natural videos recorded from primary visual cortex (*N* = 50 single cells, *N* = 25 multi unit sites), latero-intermediate visual area (*N* = 53 single cells, *N* = 33 multi unit sites), and temporal occipital cortex (*N* = 52 single cells, *N* = 26 multi unit sites). For methodological details about the experimental setup and recordings, we refer to the original paper.

#### 4.2.2 Representational dissimilarity matrices

For comparison with DNNs (see Computational modeling, Comparing neural and DNN stimulus representations), we calculated neural representational dissimilarity matrices (RDMs; Kriegeskorte et al., 2008). In short, per 16-frame video bin a neural response vector was obtained using each single and multi-unit’s average standardized firing rates. For calculating the average firing rate per 16-frame bin (spanning 533 ms), a temporal shift corresponding to the response latency which was estimated separately for each neuron or site was taken into account (see Vinken et al., 2016). This resulted in response vectors per 16-frame bin and per stimulus, which were correlated pairs-wise (Pearson *r*) in order to obtain RDMs with distances 1 − *r*. Stimulus pairs that elicit a similar neural response pattern result in a lower dissimilarity.

### 4.3 Computational modeling

#### 4.3.1 DNNs

We used three trained DNN architectures as computational models of visual processing in the primate ventral stream: AlexNet, VGG16, and VGG11-C3D. AlexNet (Krizhevsky et al., 2012) and VGG16 (Simonyan and Zisserman, 2014) were both taken from the MATLAB 2017b Deep Learning Toolbox and had been pre-trained on the ImageNet dataset (Russakovsky et al., 2015) to classify images into 1000 object categories. Both architectures have been extensively used to model ventral stream processing and object perception (Güçlü and van Gerven, 2015; Cadieu et al., 2014; Kalfas et al., 2018; Bracci et al., 2019; Kubilius et al., 2016). For the experiments involving videos we also used VGG11-C3D (called C3D in Tran et al., 2014), an architecture which is similar to VGG11 (Simonyan and Zisserman, 2014), but performs convolution and pooling across the two spatial *and* temporal dimensions (operating on 16-frame bins). This network had been pre-trained on the Sports-1M dataset (Karpathy et al., 2014) to classify videos into 487 sports categories and has previously been used to model brain responses to natural videos (Güçlü and van Gerven, 2017). All three networks consist of a sequence of convolutional and max pooling layers followed by three fully connected layers. Only AlexNet also includes local response normalization layers. Each convolutional and all except the last fully connected layers were followed by a rectifying linear activation function (ReLU). We always extracted activations from each layer before the ReLU. For each model the full set of layers is typically divided in eight layer blocks (see Table 1).

**Table 1.** Nomenclature used to refer to each architecture’s layers and the division across layer blocks (*conv* : convolutional layer, suffixes a, b, or c are used if a block contains multiple conv layers; *norm*: local response normalization; *pool* : max pooling operation; *fc*: fully connected layer).

| Layer block | AlexNet | VGG16 | VGG11-C3D |
| --- | --- | --- | --- |
| 1 | conv1<br>norm1<br>pool1 | conv1a, b<br>pool1 | conv1a<br>pool1 |
| 2 | conv2<br>norm2<br>pool2 | conv2a, b<br>pool2 | conv2a<br>pool2 |
| 3 | conv3 | conv3a, b, c<br>pool3 | conv3a, b<br>pool3 |
| 4 | conv4 | conv4a, b, c<br>pool4 | conv4a, b<br>pool4 |
| 5 | conv5<br>pool5 | conv5a, b, c<br>pool5 | conv5a, b<br>pool5 |
| 6 | fc6 | fc6 | fc6 |
| 7 | fc7 | fc7 | fc7 |
| 8 | fc8 | fc8 | fc8 |

#### 4.3.2 Feature extraction

Each network has learned a rich series of feature representations which can be accessed from every layer by obtaining unit activations from an input stimulus. We calculated these activations for every convolutional, normalization, max pooling, and fully connected layer, standardized the values across inputs and reduced the dimensionality using principal component analysis. In case of videos, activations were averaged over time within each 16-frame bin. Note that the parameters for the standardization and the transformation to principal component space were always calculated only based on the stimulus set that the animals were trained on in the behavioral experiments, thus excluding the stimuli of the test set (except for the Djurdjevic et al., 2018, experiments, where there was no separate test set for generalization). The resulting feature vectors were then used to model the behavioral task or comparison with neural representations.

#### 4.3.3 Behavioral tasks

To model the binary object discrimination tasks, we trained linear support vector machine (SVM) classifiers on the same binary tasks that the rats were trained on. The inputs for each classifier were the features extracted from a certain layer. A separate classifier was then trained for every layer on each experiment’s training set and tested for generalization on the test set to assess whether its feature representation can support the task. Each classifier was trained using the MATLAB 2017b function *fitclinear*, with the limited-memory BFGS solver and standard regularization. For tasks with one stimulus per trial (Zoccolan et al., 2009; Djurdjevic et al., 2018), the correct classification was evaluated for each object individually. For the two-alternative forced choice task in Vinken et al. (2014), a trial was considered correct if (a) the target and distractor were both on their respective correct side of the SVM boundary, (b) both were on the target side, but the target was further away from the boundary, or (c) both were on the distractor side, but the target was closer to the boundary.

In the behavioral experiments, rats were not head fixed or fixating, so the actual retinal projection of an object image could vary from trial to trial. Importantly, rats ware trained with this variability and the distributions of retinal projections during training trials should cover those during test trials. We explicitly modeled such variability during training and testing for the Zoccolan et al. (2009) experiment by varying object positions and found that this did not lead to qualitatively different results (data not shown), thus we proceeded without such variability.

We then used logistic regression to fit the positions of the stimuli in feature space relative to the SVM decision boundary onto the rat behavioral performance data obtained in the original experiment. For binary classification tasks where only one stimulus was presented on every trial (Zoccolan et al., 2009; Djurdjevic et al., 2018), we used each stimulus’ signed distance to the SVM decision boundary *d*, where positive values indicated that the stimulus was on the correct side of the boundary, and negative values indicated otherwise. For the data of Zoccolan et al. (2009), we only had percentages correct for each transformation averaged across the two objects. Thus, for a given transformation *j*, the average proportion correct *p* was predicted from the sum of the signed distances *d*_1_ and *d*_2_ for object 1 and 2 respectively:

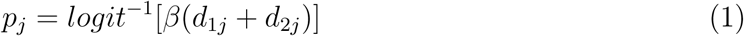

For the data of Djurdjevic et al. (2018), we had percentages correct for each object and size separately. Thus, for a given object *i* and size *j*, the average proportion correct *p* was predicted from the signed distance to the boundary *d* as follows:

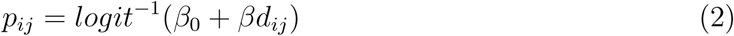

Note that we also included a bias term *β*_0_ to be able to capture the difference between good and bad performers.

In the two-alternative forced choice task (Vinken et al., 2014) a target and distractor were presented simultaneously and could both be directly compared by the rats. To model the comparison between target and distractor, we used the distance between target and distractor along a readout dimension orthogonal to the SVM boundary. To calculate this distance, we took the signed distance to the decision boundary (positive values for the “rat” side of the boundary, negative values otherwise) of the target stimulus, minus the signed distance to the decision boundary of the distractor stimulus (also positive values for the “rat” side of the boundary, negative values otherwise). Thus, for a given target-distractor (16-frame bin) pair *i*, the average proportion correct *p* was predicted from the difference between the signed distances for the target *d*_*t*_ and distractor *d*_*d*_ stimuli:

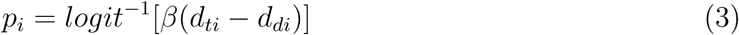

This approach allowed us to estimate how much transformation or stimulus-level variance was explained by the relative positions of stimuli in a layer’s feature space.

#### 4.3.4 Comparing neural and DNN stimulus representations

In order to estimate how closely the representational geometry of each DNN layer matched that of visual areas along the lateral stream, we calculated RDMs based on DNN features. As for the neural data, the stimulus feature vectors (one vector for each 16-frame video bin) were correlated pairs-wise in order to obtain an RDM for each DNN layer. Again, stimulus pairs that share a similar representation across features in a layer result in a lower dissimilarity. We then quantified the correspondence between neural and DNN RDMs by calculating the Spearman correlation between off-diagonal upper halves of the matrices. We normalized the correlations between neural and DNN RDMs by dividing by each area’s noise ceiling. To estimate the noise ceiling we split the trials per movie in two halves and computed the Spearman correlation between the two resulting neural RDMs (one from each split half). The noise ceiling was the Spearman-Brown-corrected average (across 1000 random splits) split-half correlation.

## 5 Acknowledgements

This work was supported by Research Foundation Flanders, Belgium (PhD and postdoctoral fellowships of K.V.); by KU Leuven Research Council (C14/16/031); by Excellence of Science (EOS) grant HUMVISCAT. We thank Davide Zoccolan and Thomas P. O’Connell for helpful comments on this work.

